# SYBR green one-step qRT-PCR for the detection of SARS-CoV-2 RNA in saliva

**DOI:** 10.1101/2020.05.29.109702

**Authors:** DR Ganguly, S Rottet, S Yee, WY Hee, AB Smith, NC Khin, AA Millar, AM Fahrer

## Abstract

We describe our efforts at developing a one-step quantitative reverse-transcription (qRT)-PCR protocol to detect severe acute respiratory syndrome coronavirus 2 (SARS-CoV-2) RNA directly from saliva samples, without RNA purification. We find that both heat and the presence of saliva impairs the ability to detect synthetic SARS-CoV-2 RNA. Buffer composition (for saliva dilution) was also crucial to effective PCR detection. Using the SG2 primer pair, designed by Sigma-Aldrich, we were able to detect the equivalent of 1.7×10^6^ viral copies per mL of saliva after heat inactivation; approximately equivalent to the median viral load in symptomatic patients. This would make our assay potentially useful for rapid detection of high-shedding infected individuals. We also provide a comparison of the PCR efficiency and specificity, which varied considerably, across 9 reported primer pairs for SARS-CoV-2 detection. Primer pairs SG2 and CCDC-N showed highest specificity and PCR efficiency. Finally, we provide an alternate primer pair to use as a positive control for human RNA detection in SARS-CoV-2 assays, as we found that the widely used US CDC primers (targeting human *RPP30*) do not span an exon-exon junction and therefore does not provide an adequate control for the reverse transcription reaction.

## Introduction

The ongoing SARS-CoV-2 pandemic continues to rapidly spread globally, reaching in excess of 5.2 million cases and 330,000 deaths, causing substantial socio-economic impacts (UNSDG 2020; Nicola et al. 2020; WHO 2020a). In order to facilitate: (I) minimising and tracing the spread of SARS-CoV-2, and (II) an easing of lockdown measures; we require information on the extent of community transmission, particularly of asymptomatic cases (Prather et al. 2020). This necessitates accessible, high-throughput, and cost-effective methods of SARS-CoV-2 detection.

Quantitative reverse-transcription (qRT)-PCR has become a critical tool for detecting SARS-CoV-2, by amplifying virus-derived RNA, due to its increased sensitivity and fast processing time (Esbin et al. 2020). However, this has been limited by the requirement for labor-intensive RNA extractions in order to enrich viral RNA. We propose an alternative method that bypasses RNA extractions by performing one-step qRT-PCR directly on saliva. Importantly, saliva samples have been highlighted as a useful diagnostic tool for SARS-CoV-2 detection (Azzi et al. 2020) and qRT-PCR on crude saliva samples has been used to detect Zika virus (Li et al. 2019). We also propose the use of SYBR green-based detection to complement the multiplexed hydrolysis probe-based detection of widely used kits. The relative simplicity and cost of SYBR green detection is particularly relevant for facilitating mass testing of SARS-CoV-2 in poorer countries. Indeed, SYBR green, *Taq* polymerase, and reverse transcriptase can all be made relatively cheaply and there is a growing community effort for open source protocol development (e.g. BEARmix: https://gitlab.com/tjian-darzacq-lab/bearmix).

The use of SYBR green also allows for the diagnosis of non-specific amplification via melt curve analysis (T_m_ calling). Indeed, non-specific amplification has been a problem for hydrolysis probe-based kits, likely the result of sequence similarity to RNA from other commonly found pathogens including viruses (Vogels et al. 2020; Rahman et al. 2020). For example, early versions of the US CDC kits were prone to false positives when testing for SARS-CoV-2 because the primers bound to other milder strains of coronavirus (Satyanarayana 2020; Cohen 2020). Therefore, the ability to distinguish non-specific amplification is important for reliable diagnosis. Finally, the internal control primers from the US CDC kit [targeting human *RPP30* (Vogels et al. 2020)] do not span an exon-exon junction thus allowing amplification of genomic DNA instead of specifically detecting RNA-derived cDNA. This means it does not provide a reliable positive control for successful reverse transcription. We also tested the combined use of a sample buffer containing Tween20 and heat treatment for impediments on PCR chemistry so they could be considered for virus inactivation (Roberts et al. 2009; Darnell et al. 2004).

## Method

Human RNA was purified from blood using TRIzol (Invitrogen) as per the manufacturer’s instructions. Synthetic SARS-CoV-2 RNA Control 1 (Twist Bioscience, SKU:102019) was used to represent viral RNA. One-step qRT-PCR was performed using the Power SYBR one-step kit (Applied Biosystems) to amplify RNA-derived cDNA in 10 µL reactions on a LightCycler480 Instrument II (Roche, LC480).

The following describes our protocol use to obtain the results herein:

1. Saliva samples were collected in a 50 mL Falcon tube.
2. Samples were diluted 1:1, using a reverse pipetting technique with wide-end or cut pipette tips to overcome saliva viscosity, with sample buffer (TE-T: 10 mM Tris pH 6.5, 1 mM EDTA, 1% Tween20) in a clean PCR tube then heated at 95 °C for 5 mins.
3. Heat-treated samples were further diluted to a final dilution ranging from 1/4 to 1/128.
4. A one-step RT-PCR master mix was prepared containing Power SYBR Green RT-PCR Mix (2×, 5 µL per reaction), 125× RT enzyme mix (0.08 µL per reaction), and appropriate primer pairs (0.15 µL of 10 µM stock for each primer per reaction to obtain final concentration of 150 nM), according to the manufacturer’s instructions for a 10 uL reaction volume. Primers used are listed in Table 1.
5. 5.4 µL of the master mix was combined with 4.6 µL of sample on a 384-well skirted PCR plate (Roche style, BioTools). Samples include: (I) saliva diluted in sample buffer, (II) purified human RNA diluted in nuclease-free water, (III) synthetic SARS-CoV-2 RNA diluted in nuclease-free water, or (IV) no-template control (NTC: nuclease-free water or sample buffer as appropriate).
6. All samples were run in either technical duplicate or triplicate reactions.
7. The plate was sealed with an adhesive optically transparent PCR seal (Integrated Sciences).
8. The LC480 was run according to the program outlined in Table 2 based on the manufacturer’s instructions.
9. The LC480 Instrument Software (v1.5.0) was used to calculate threshold cycle (Ct) values from raw fluorescence. First, T_m_ calling was performed on all reactions to exclude:
  a. Reactions with products matching non-specific amplification as reflected from the negative control (no template), or
  b. Reactions not demonstrating a single clear product matching the product amplified from the positive control (SARS-CoV-2 RNA Control 1).
10. Amplified products from each primer pair were inspected by gel electrophoresis (1.5% agarose) to match the T_m_ calling results to expected fragment size (Figure 4 A-B).
11. The *2nd derivative method* was used for determining Ct values (based on polynomial regression performed by the LightCycler480 Instrument Software using the high confidence algorithm). Each reaction, and its corresponding Ct value, was considered individually.
12. PCR efficiency was estimated using two separate methods:
  a. PCR efficiency for each primer pair was estimated using LinReg on positive control reactions demonstrating a single expected product as determined in step 8 (Ruijter et al. 2009). Additionally, PCR efficiencies were only considered if reactions passed LinReg sample and quality checks, for example reactions displaying minimal noise, no baseline error, and appropriate amplification and plateau phases. In some cases PCR efficiencies could not be determined by LinReg (“n.d.”), for example due to variable background fluorescence and/or minimal amplification.
  b. For selected primer pairs, PCR efficiency was also estimated using a dilution series of Twist Synthetic SARS-CoV-2 RNA Control 1 (or purified human RNA for *GAPDH*). A standard curve was constructed in GraphPad Prism (Ct values plotted against Log_10_ Concentration) and its slope was used to determine PCR efficiency with the formula: E = [-1+10^−1/slope^] x 100.

**Figure 1.**
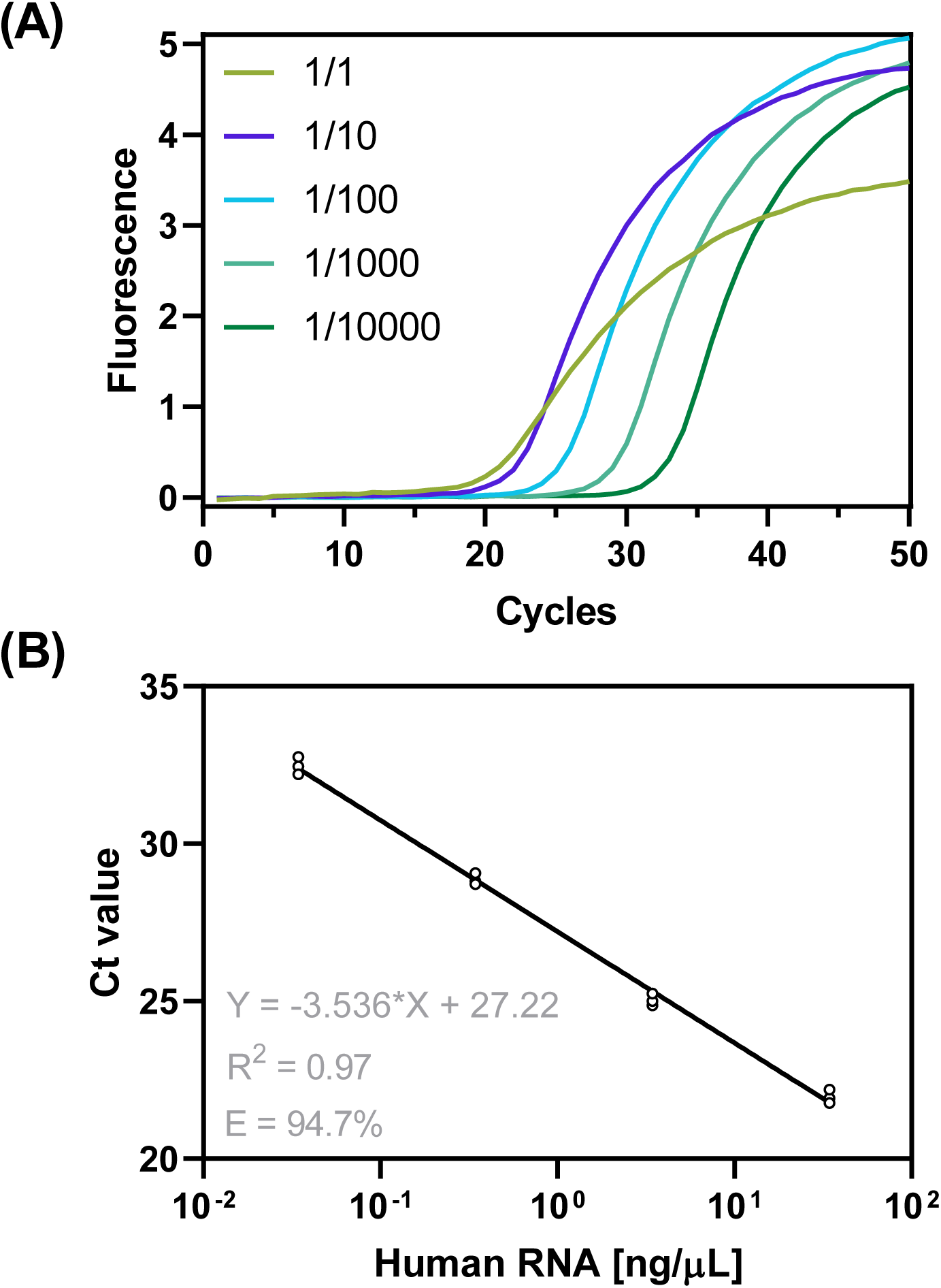
*Amplification of GAPDH* from purified human RNA. **(A)** Raw fluorescence curves obtained using *GAPDH* primers on human RNA dilutions ranging from 1x (344.4 ng/µL) to 1/10,000 (0.034 ng/µL). Data represent means from technical replicates (n=3). (**B**) Linear regression of Ct against RNA concentration (ng/µL) to calculate PCR efficiency of *GAPDH* primers (E=94.7%, n=3). The 1x sample was excluded from the regression.

**Figure 2.**
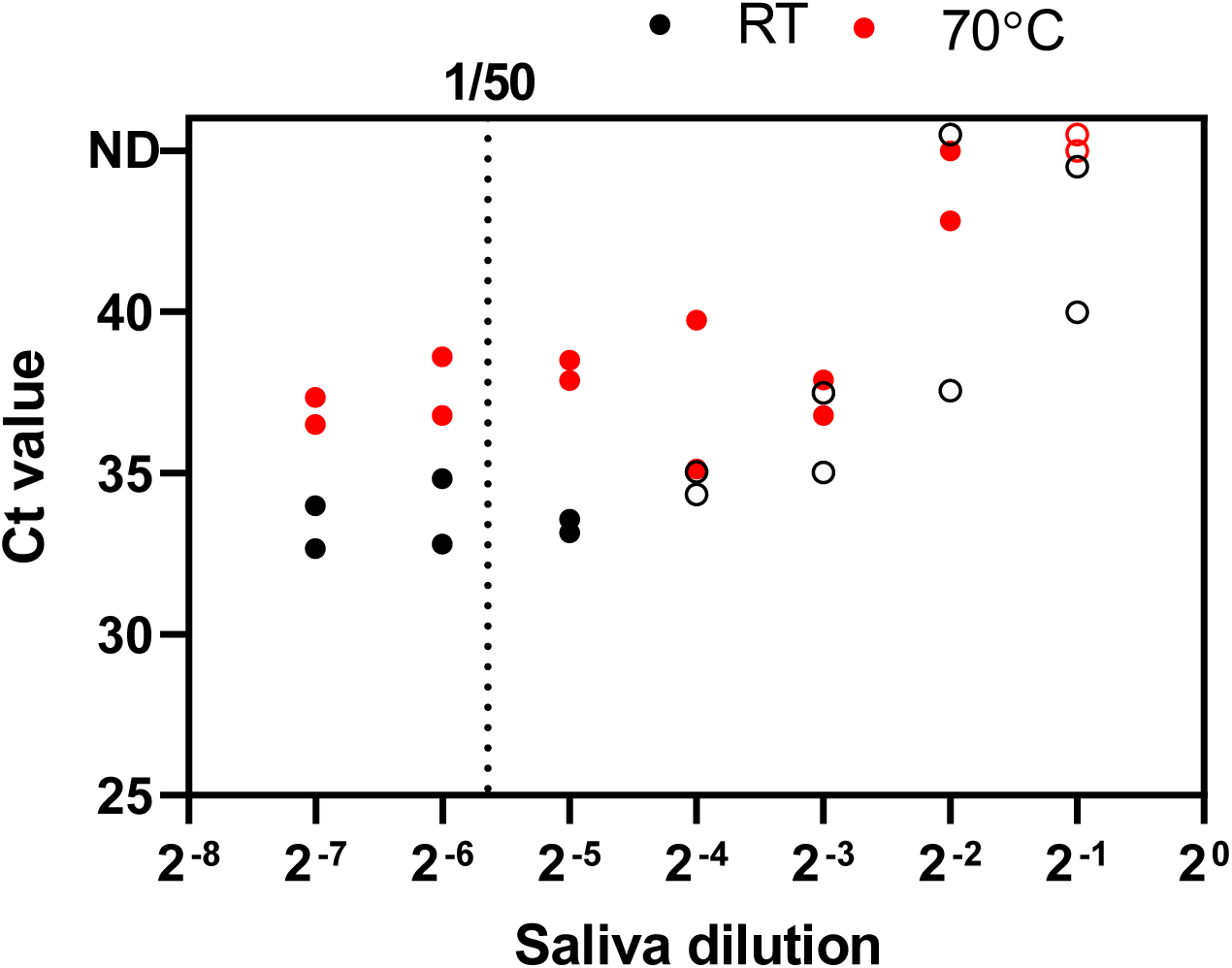
*GAPDH* amplification across saliva dilution series. Ct values for *GAPDH* amplification from saliva diluted in TE-T (1/2, 1/4, 1/8, 1/16, 1/32, 1/64, 1/128). Diluted saliva was then either treated at 70°C for 5 min (*red circles*) or left at room temperature for 5 min (*black circles*). Points denote individual biological replicates (n=2). Non-specific amplicons (based on T_m_ calling) are depicted as *open circles*.

**Figure 3.**
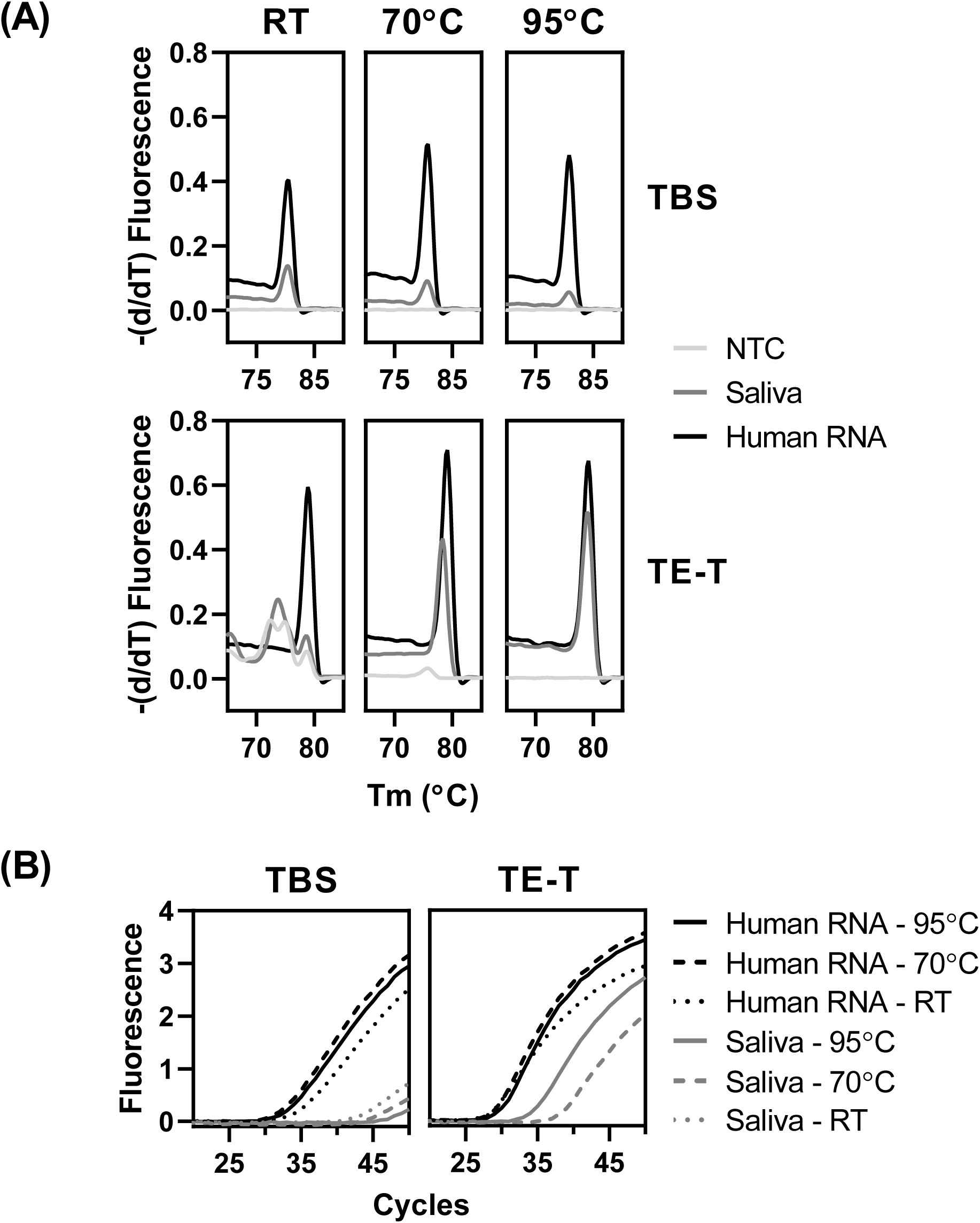
Testing buffer composition and heat treatments on *GAPDH* amplification. (**A**) Plots showing the absolute value of the first derivative of raw fluorescence values obtained during the melt curve analysis for *GAPDH* primers. (**B**) Plots showing raw fluorescence curves captured during *GAPDH* amplification. Saliva, human RNA and NTC were resuspended in either TBS (20 mM Tris pH 7.2, 150 mM NaCl) or TE-T (10 mM Tris pH 6.5, 1 mM EDTA, 1% Tween20), and heat-treated (70 or 95°C for 5 min) or kept at room temperature (RT). Plots represent means from biological replicates (n=2). Note, saliva at RT in TE-T buffer amplified non-specific products so was excluded.

**Figure 4.**
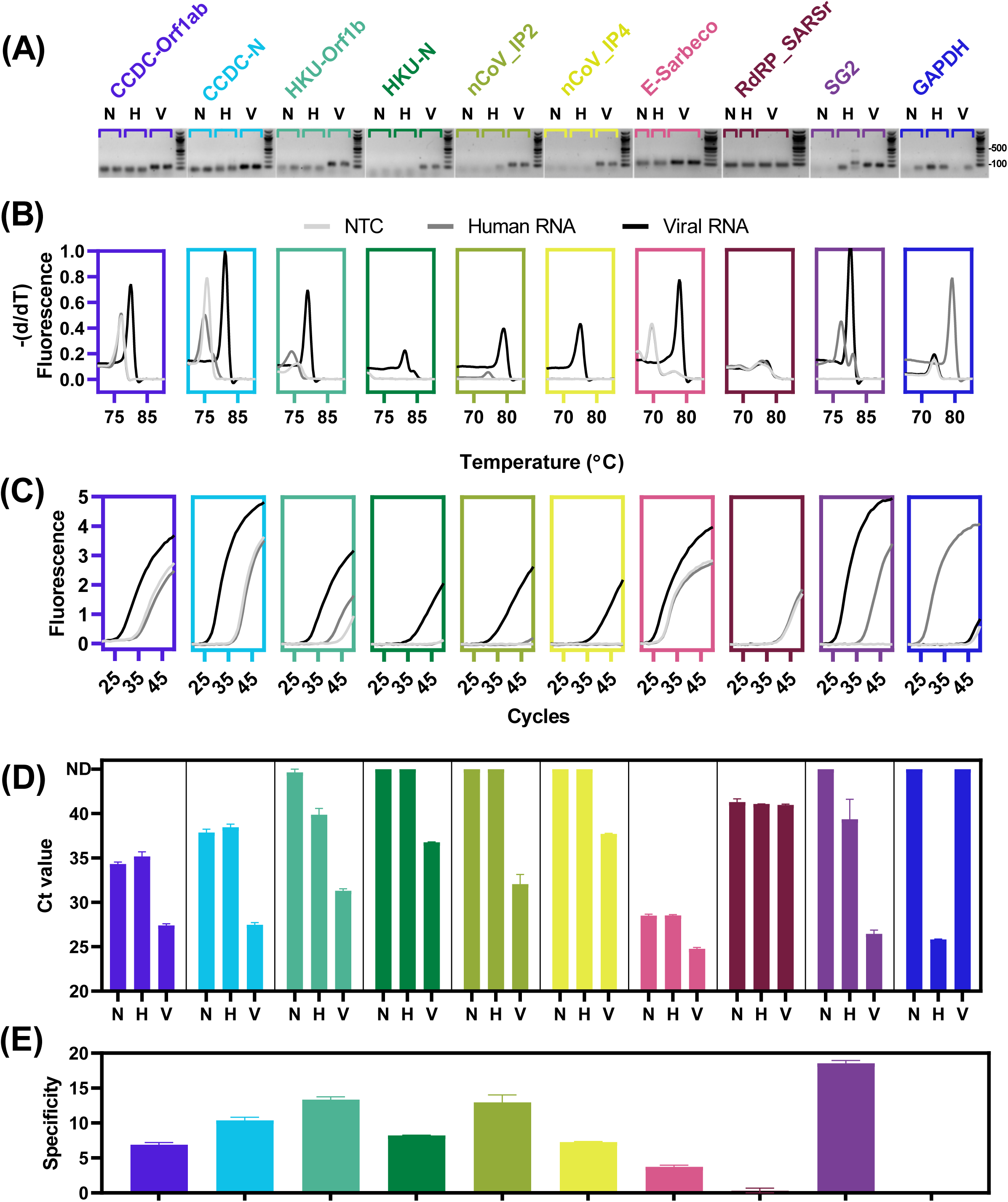
Primer screening for SARS-CoV-2 detection. (**A**) Gel electrophoresis of individual qRT-PCR products. (**B**) Plots showing the absolute value of the first derivative of raw fluorescence values obtained during the melt curve analysis for each primer pair. (**C**) Plots showing raw fluorescence curves captured during amplification of each primer pair. (**B**) and (**C**) Plots represent means from biological replicates (n=2). (**D**) Ct values obtained for the amplification of each primer pair tested for SARS-CoV-2 detection. (**E**) Calculated specificity for each primer (Ct_NTC_–Ct_Viral_). Data represent the means of biological replicates (n=2). Error bars denote SEM. Abbreviations: N or NTC, no template control; H, purified human RNA; V, viral RNA (Synthetic RNA Control 1).

## Results

We first established a workflow for one-step qRT-PCR by amplifying *GAPDH* from a dilution series of purified human RNA (1x = 344.4 ng/µL; Figure 1 A-B). The reverse primer spans an exon-exon junction in *GAPDH* (exons 2 and 3), thus minimising the risk of amplifying genomic DNA (McIntyre et al. 2012). As expected, a single product was observed and Ct values demonstrated a log-linear relationship to RNA concentration. The abnormal fluorescence curve from in the 1x sample (not included in regression) suggested the possibility of a saturated reverse-transcription reaction resulting in a less pronounced amplification phase.

**Table 1.**
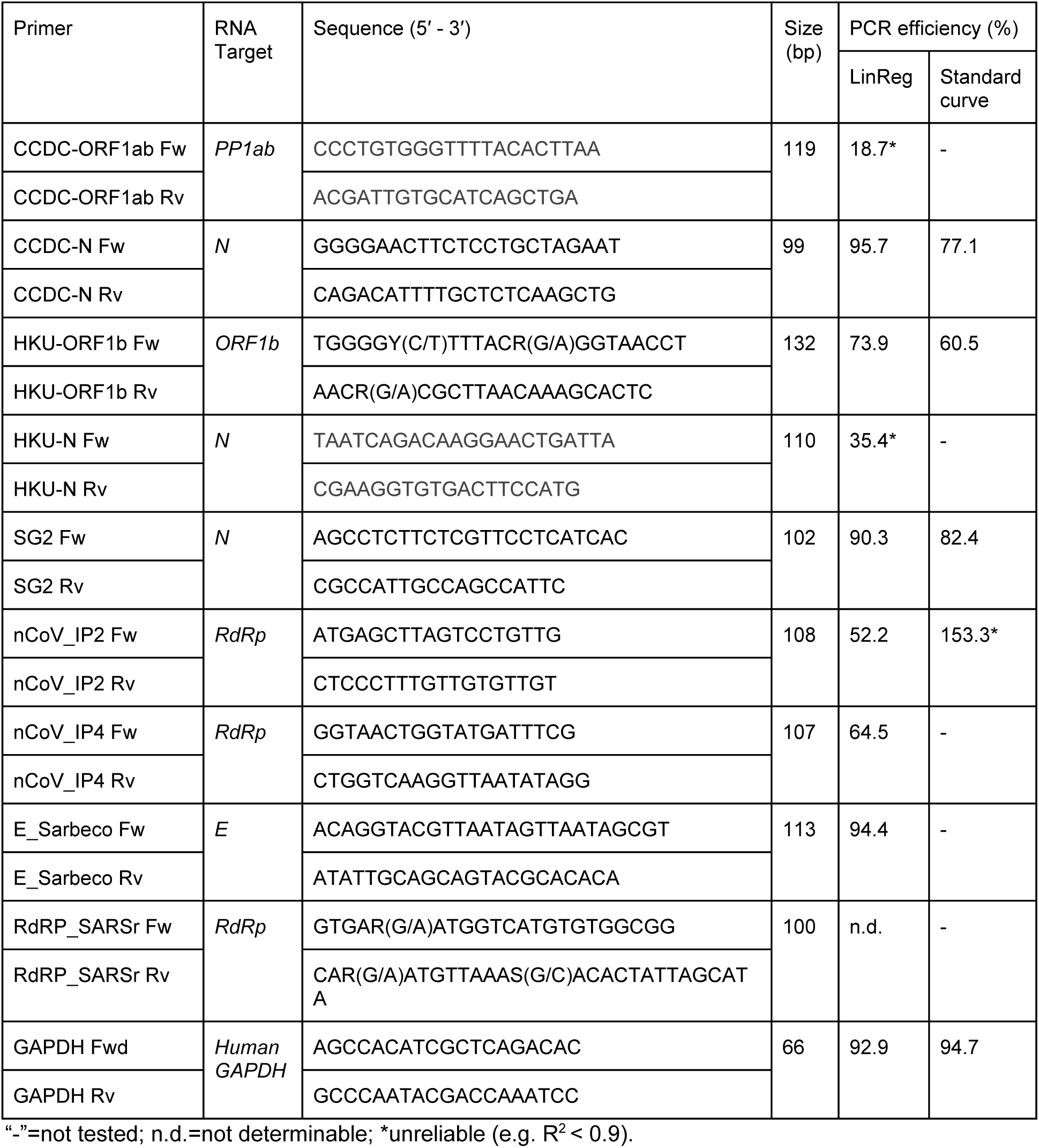
Primer information for SARS-CoV-2 qRT-PCR assays

**Table 2.** LightCycler480 Run Protocol

| Stage | Step | Temperature (°C) | Hold time (mm:ss) | Acquisition (483-533 nm) | Ramp Rate (°C/s) |
| --- | --- | --- | --- | --- | --- |
| Holding | Reverse Transcription | 48 | 30:00 | None | 4.8 |
| Holding | Activation of polymerase | 95 | 10:00 | None | 4.8 |
| Cycling (40 - 50x) | Denature | 95 | 00:15 | None | 4.8 |
|  | Anneal and Extend | 60 | 1:00 | Single | 2.5 |
| Melt curve | Denature | 95 | 00:30 | None | 4.8 |
|  | Anneal | 60 | 00:30 | None | 2.5 |
|  | Denature | 95 |  | Continuous | 0.11 |

We next amplified *GAPDH* from a series of diluted saliva samples without RNA purification (Figure 2). We checked for specific amplification by comparing product T_m_ based on purified human RNA. Unfortunately, high concentrations of saliva inhibited the reaction as seen by open circles in 2^-1^ - 2^-4^ dilutions. As we diluted the saliva, we were able to more confidently amplify *GAPDH* suggesting that some saliva-derived material may be interfering with the reaction. We also tested the effect of a moderate heat treatment (70°C for 5 minutes). This was done to simulate heat inactivation, which would be performed on saliva from potentially infectious samples. Interestingly, whilst Ct values increased after heat-treatment, heat-treatment improved the ability to amplify the expected product from *GAPDH* in less diluted saliva.

We next tested whether the use of various sample buffers and heat treatments impaired the one-step RT-PCR reaction. This included the use of Tween20 and a 5 minute 95°C heat treatment, which is advised for inactivating viral particles (Roberts et al. 2009; Darnell et al. 2004). Three buffers were trialled: PBS (pH 7.4, 10 mM Na_2_HPO_4_, 2 mM KH_2_PO_4_, 2.7 mM KCl, 140 mM NaCl), TBS (20 mM Tris pH 7.2, 150 mM NaCl), and TE-T (10 mM Tris pH 6.5, 1 mM EDTA, 1% Tween20) alongside two heat treatments (70 or 95°C) to observe whether *GAPDH* amplification was impeded. The use of PBS led to no amplification regardless of sample or treatment. On the other hand, TBS and TE-T could amplify *GAPDH* from human RNA for all conditions (Figure 3 A-B; note that TE-T shifted the expected T_m_ by approximately 3 degrees compared to TBS). TE-T was clearly superior to TBS in amplifying *GAPDH* from saliva. Finally, the use of a 5 minute 95°C heat treatment led to improved *GAPDH* amplification from saliva (Figure 3 B).

Using these conditions, we next sought to test a range of primers designed for SARS-CoV-2 detection. As we were using a SYBR green-based method, we sought to minimize the possibility of non-specific amplification. The combination of NCBI Primer-BLAST (Ye et al. 2012) and Nucleotide BLAST was used to screen a range of primers, designed to detect SARS-CoV-2 RNA, based on the following criteria:

1. Primer-BLAST: no products <1000 bp from human mRNA and genomic DNA
2. Nucleotide BLAST: no, or minimal, matches to human mRNA and genomic DNA
3. For *GAPDH*, one of the primers had to span an exon-exon junction to minimise the risk of amplifying genomic DNA.

From this process we selected the following primers: CCDC-N and CCDC-ORF1ab (China CDC), HKU-ORF1b and HKU-N (University of Hong Kong), SG2 (Sigma-Aldrich), nCoV_IP2 and nCoV_IP4 (Institut Pasteur), and E_Sarbeco and RdRP_SARSr (Charité) [Table 1 (WHO 2020b)].

We tested the ability, of the selected primers, to amplify synthetic SARS-CoV-2 RNA. Figure 4 shows our results for nine SARS-CoV-2 primer pairs, including gel electrophoresis to confirm correct amplification (Figure 4 A), melt curve analysis (Figure 4 B), and SYBR-green fluorescence (Figure 4 C). Subsequently, we calculated Ct values from the raw fluorescence curves for all reactions (Figure 4 D). Using these calculated Ct values, we determined the specificity for each primer with which to compare performance (Ct_NTC_–Ct_Virus_; Figure 4 E). Based on this, we selected four primer pairs: CCDC-N, HKU-ORF1b, nCoV_IP2, and SG2; for further analysis.

We next performed qRT-PCR on a dilution series of synthetic SARS-CoV-2 RNA to determine a limit of detection (LoD) per select primer pair and included saliva, human RNA, and NTC as negative controls (Figure 5). Based on the ability to amplify the correct product, based on T_m_ calling, and observing a log-linear relationship between Ct value and viral RNA concentration. We estimate the LoD for CCDC-N (64–256 copies per reaction), HKU-ORF1b (256-1024 copies per reaction), and SG2 (16–64 copies per reaction). We could not determine a LoD for nCoV_IP2, although this primer pair appeared less prone to amplifying non-specific products.

**Figure 5.**
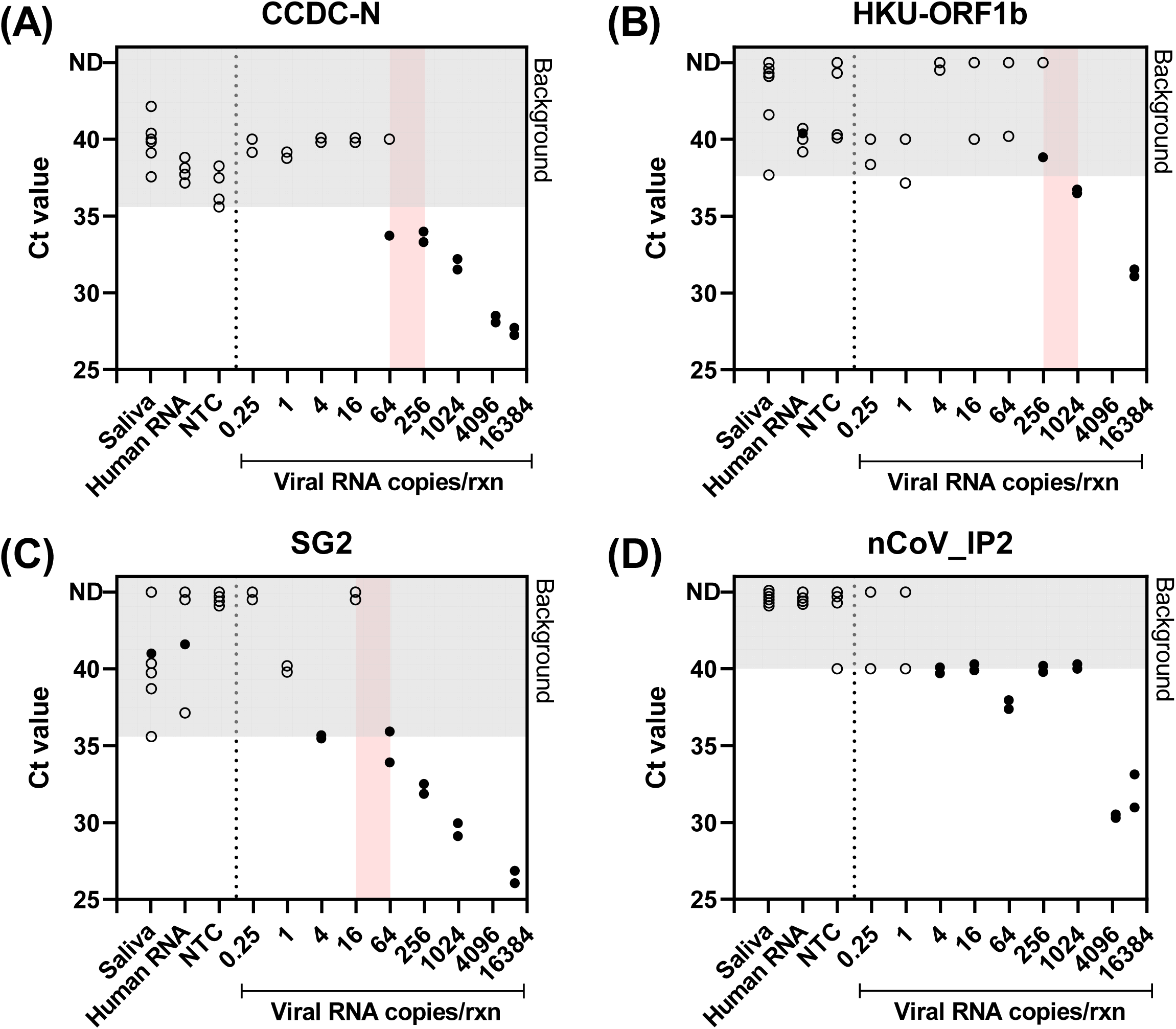
Estimating the limit of detection for selected primer pairs Scatterplots representing Ct values for select primer pairs across a dilution series of synthetic SARS-CoV-2 RNA. Points reflect individual data points for reactions with saliva (dilution=1/50, n=6), purified human RNA (quantity=1.62 ng, n=4), NTC (n=4), and viral RNA dilutions (n=2). *Open circles* denote reactions producing non-specific amplicons based on Tm calling. *Closed circles* denote reactions producing an amplicon matching the expected product. *Grey shaded area* represents the background amplification and is defined by the lowest Ct value from the negative controls. Approximate LoD is represented by the *red shaded area*.

Lastly, we tested whether saliva interfered with the detection of SARS-CoV-2 RNA. To do this, we spiked-in two quantities of SARS-CoV-2 RNA (100 or 1000 copies per reaction) into a saliva dilution series. We picked dilutions ranging from 1/8 to 1/64 to identify the minimal dilution allowing unimpeded detection using our most sensitive primer pairs: SG2 and CCDC-N (Figure 6). Unfortunately, mixing with saliva resulted in reduced sensitivity (higher Ct values). Furthermore, heat treatment abolished the sensitivity of CCDC-N at both RNA concentrations and for SG2 on 100 copies per reaction. This is, in part, likely driven by heat-induced RNA degradation (see Figure 6, right, “100 copies of Viral RNA”). Promisingly, primer pair SG2 could detect 1000 copies of SARS-CoV-2 RNA in all saliva dilutions after heat treatment (Figure 6, top left panel).

**Figure 6.**
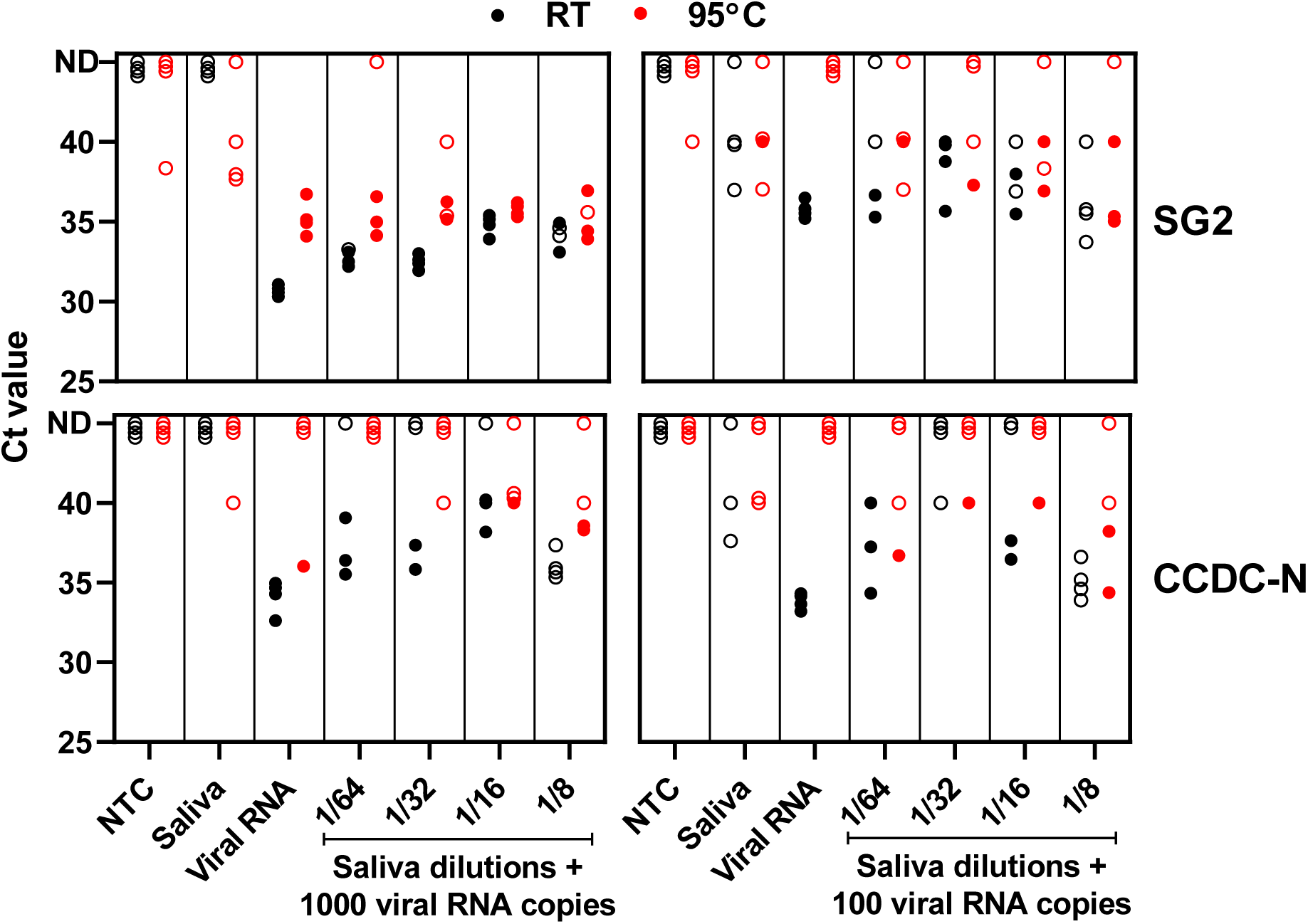
Influence of saliva and heat treatment on SARS-CoV-2 RNA detection Ct values determined using primer pairs SG2 (*top*) and CCDC-N (*bottom*) on a saliva dilution series with and without heat treatment (5 minutes 95°C) containing 1000 (*left*) or 100 (*right*) copies per reaction of SARS-CoV-2 RNA. *Closed circles*, correct T_m_; *open circles*, non-specific T_m_; *black circles*, room temperature (RT) for 5 min; *red circles*, 95°C for 5 min; NTC, no template control; S, saliva only. Saliva dilutions range from 1/8 to 1/64. Points represent individual data points obtained from biological (n=2) and technical (n=2) replication. Overlapping points were offset for clarity.

## Discussion

From these results, we describe a SYBR green-based approach for the detection of SARS-CoV-2 directly from saliva samples. Performing a one-step qRT-PCR assay directly on saliva provides several advantages, including relative ease of self-collection requiring no trained personnel and less PPE, minimal patient discomfort, skipping laborious and expensive RNA extractions, and reducing demand on swabs (which are in global shortage) (Azzi et al. 2020; Xu et al. 2020). The SYBR green-based methodology also offers the ability to diagnose non-specific amplification, which is not possible using hydrolysis-based primer-probes, and reduces demand on commercial kits, which are in short supply (thereby increasing testing capacity). We estimate the cost of these reactions to be in the range of AUD $4-8 per sample. Aspects of this method could be combined with community-driven open source protocols (for example, *BEARmix*: https://gitlab.com/tjian-darzacq-lab/bearmix) to achieve even lower costs while addressing technical limitations, such as appropriate methods for virus inactivation as discussed below.

Despite promising progress, we must highlight the critical need for validation against patient samples alongside clinical certification in order for results to be considered diagnostic. While we attempted to mimic patient samples by spiking synthetic RNA into non-infectious saliva, this cannot replace validation using patient saliva. We propose the combined use of Tween20 and heat treatment (95 °C for 5 mins) for viral inactivation (Roberts et al. 2009; Darnell et al. 2004), and for increased release of viral RNA from patient saliva. However, this needs to be validated with samples from patients with known SARS-CoV-2 infection. We therefore report a theoretical limit of detection based on reactions using synthetic virus RNA while controlling for non-specific amplification.

Primer pairs CCDC-N and SG2 demonstrated the best specificity (Figure 4 E) and sensitivity (Figure 5 A, C) towards the synthetic SARS-CoV-2 RNA. Based on the amplification of synthetic SARS-CoV-2 RNA in serial dilutions (no saliva), we estimate the LoD for CCDC-N and SG2 primer pairs to be 64-256 and 16-64 molecules per reaction, respectively. Therefore, we propose these primers to be best suited for SARS-CoV-2 screening reactions. However, whilst an LoD could not be determined for nCoV_IP2, this primer pair showed negligible non-specific amplification (leading to higher specificity) and, therefore, would be useful to confirm ambiguous samples. The amplification efficiency observed here performs comparably with other primer-probe based detection methods on synthetic SARS-CoV-2 RNA (Vogels et al. 2020). Promisingly, despite different methodology and source RNA, we arrive at similar conclusions regarding the superiority of nCoV_IP2 (or RdRp IP2) and CCDC-N primers, whereas RdRP_SARSr and E_Sarbeco performed poorly (Etievant et al. 2020; Vogels et al. 2020).

We uncover a promising new candidate in primer pair SG2 designed by Sigma-Aldrich for research use (https://www.sigmaaldrich.com/covid-19.html). With these primers, we estimate a LoD of 16 - 64 molecules per reaction, which translates to between 1.6×10^4^ – 6.4×10^4^ copies per mL in the original sample (Table 3). The viral load in self-collected saliva samples has been estimated to range from 9.9×10^2^ – 1.2×10^8^ (median = 3.3×10^6^) copies per mL (To et al. 2020b). The median viral load in posterior oropharyngeal saliva samples has been estimated as 1.6×10^5^ copies per mL (To et al. 2020a). However, these measures of viral load were from patients showing disease symptoms. It remains unclear what the viral load in asymptomatic cases are, though there is some limited evidence for a correlation between symptom severity and the amount of viral particles detected (Liu et al. 2020). Thus, if purifying RNA from saliva, the SG2 primer pair will be able to detect the majority of infected individuals (Table 3, SG2 LoD = 1.6×10^4^ - 6.4×10^4^ copies per mL).

**Table 3.**
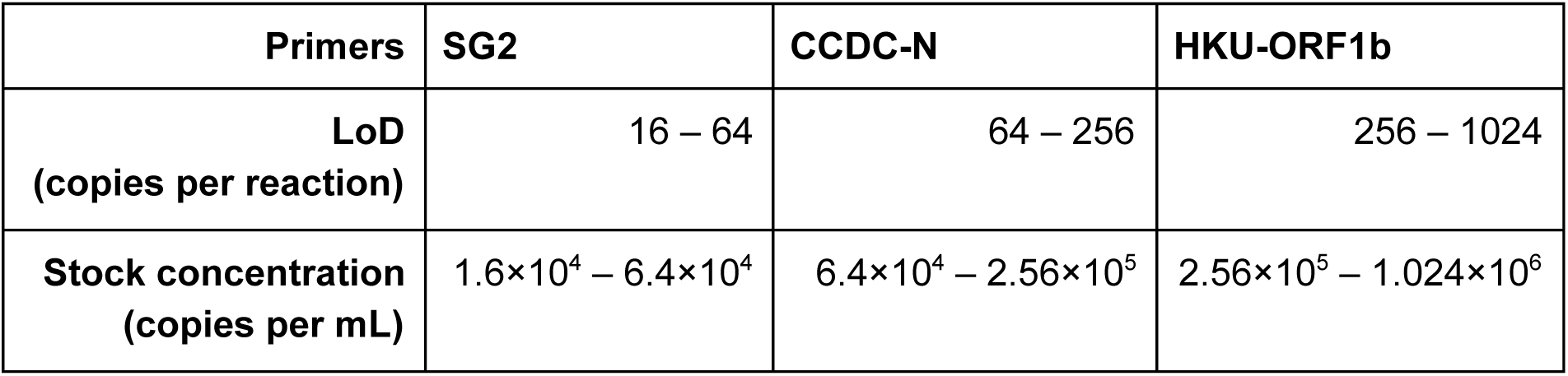
Estimated LoD for highest performing primer pairs amplifying synthetic SARS-CoV-2 RNA

| Primers | SG2 | CCDC-N | HKU-ORF1b |
| --- | --- | --- | --- |
| **LoD (copies per reaction)** | 16 – 64 | 64 – 256 | 256 – 1024 |
| **Stock concentration (copies per mL)** | $1.6 \times 10^4 - 6.4 \times 10^4$ | $6.4 \times 10^4 - 2.56 \times 10^5$ | $2.56 \times 10^5 - 1.024 \times 10^6$ |

SARS-CoV2-RNA could be detected directly from saliva (mimicking patient samples) without purification. Whilst this was successful for samples kept at room temperature, our heat treatment led to reduced sensitivity, especially for the CCDC-N primer pair. Nonetheless, primer pair SG2 was still able to detect 1,000 RNA copies per reaction in saliva diluted 1/8 (of which 4.6 µL is added to the reaction), translating to approximately 1.7×10^6^ copies per mL after heat-treatment, which is still within the viral load range. The heat-induced reduction in sensitivity against SARS-CoV-2 has also been reported independently (Pan et al. 2020). The reduced sensitivity is likely the result of: (I) increased spontaneous RNA cleavage at higher temperatures, and (II) enzymatic degradation by salivary ribonucleases post-heat treatment, which may also denature protective proteins or RNA secondary structures (Brisco and Morley 2012; Emilsson et al. 2003). This degradation may be circumvented by the addition of RNA stabilisers, such as 6-8% formamide (Yasukawa et al. 2010), prior to heat treatment or with an alternate buffer with greater pH stability at higher temperatures (Sullivan et al. 2020; Reineke et al. 2011). However, caution is required as buffer composition drastically affects RT-PCR performance as highlighted by no amplification with PBS, although it has been shown to be a suitable medium for transporting samples (Rodino et al. 2020). Alternative methods for viral inactivation could be considered, such as the addition of biocidal agents, proteases, stronger detergents, or reducing agents (Brittany S. Mertens 2015; Pfaender et al. 2015; Carver and Seto 1968; Leung et al. 2017; Kampf et al. 2020). This would require subsequent inactivation and/or dilution to prevent the impediment of PCR enzymes. It may also be possible to apply ultraviolet radiation to saliva samples or temporarily alter the pH, however, it may be challenging to find a treatment leading to viral inactivation without complete RNA damage (Darnell et al. 2004; Lemire et al. 2016; Beck et al. 2015). Alternatively, RNA extractions could be performed on saliva for which clinically-relevant methods now exist, however, this would still add considerable costs and labour (Pandit et al. 2013; Sullivan et al. 2020). Methods for one-step RNA extractions may provide a compromise between direct qRT-PCR and complete RNA extraction (Sentmanat et al. 2020).

We hope our results will be helpful to other laboratories investigating the possibility of one-step qRT-PCR for SARS-CoV-2 testing from saliva samples. Our comparison of various reported primer pairs is also likely to be of use to laboratories around the world. In particular, we hope this information provides a platform for increased accessibility to promote SARS-CoV-2 testing, especially in poorer countries. Finally, we propose the use of the *GAPDH* primer pair, used here, as the human RNA positive control. The reverse primer spans an exon-exon junction unlike the widely used US CDC primers.

## Author Contributions

Conceptualization, A.F. and A.M.; methodology, A.S. and D.G.; software, N/A; validation, S.Y., W.H., and N.K.; formal analysis, S.R. and D.G.; investigation, S.Y., W.H., S.R., A.S., N.K. and D.G.; resources, A.M. and A.F.; data curation, S.R., S.Y. and D.G.; writing—original draft preparation, D.G. and A.F.; writing—review and editing, D.G., S.R., and S.Y.; visualization, S.R.; supervision, A.F and A.M.; project administration, A.F. and D.G.; funding acquisition, A.F and A.M. All authors have read and agreed to the published version of the manuscript.

## Acknowledgements

We would like to acknowledge the helpful and critical feedback with Tamás Fischer, Rippei Hayashi, Thomas Tapmeier, and Anselm Enders. Primers targeting *GAPDH* were kindly provided by Thomas Tapmeier and Tamás Fischer.

This research was funded by the Research School of Biology at The Australian National University. D.G., A.S., and N.K. were supported by the ARC Centre of Excellence in Plant Energy Biology (CE140100008). D.G. was supported by the CSIRO Synthetic Biology Future Science Platform.

The authors declare no conflict of interest.

## References

Azzi L, Carcano G, Gianfagna F, Grossi P, Gasperina DD, Genoni A, Fasano M, Sessa F, Tettamanti L, Carinci F, et al. 2020. Saliva is a reliable tool to detect SARS-CoV-2. J Infect. http://dx.doi.org/10.1016/j.jinf.2020.04.005.

Beck SE, Rodriguez RA, Hawkins MA, Hargy TM, Larason TC, Linden KG. 2015. Comparison of UV-Induced Inactivation and RNA Damage in MS2 Phage across the Germicidal UV Spectrum. Appl Environ Microbiol 82: 1468–1474.

Brisco MJ, Morley AA. 2012. Quantification of RNA integrity and its use for measurement of transcript number. Nucleic Acids Res 40: e144–e144.

Brittany S. Mertens ODV. 2015. Characterization and Control of Surfactant-Mediated Norovirus Interactions. Soft Matter 11: 8621.

Carver DH, Seto DS. 1968. Viral inactivation by disulfide bond reducing agents. J Virol 2: 1482–1484.

Cohen J. 2020. The United States badly bungled coronavirus testing—but things may soon improve. Science, February 28 https://www.sciencemag.org/news/2020/02/united-states-badly-bungled-coronavirus-testing-things-may-soon-improve.

Darnell MER, Subbarao K, Feinstone SM, Taylor DR. 2004. Inactivation of the coronavirus that induces severe acute respiratory syndrome, SARS-CoV. J Virol Methods 121: 85–91.

Emilsson GM, Nakamura S, Roth A, Breaker RR. 2003. Ribozyme speed limits. RNA 9: 907–918.

Esbin MN, Whitney ON, Chong S, Maurer A, Darzacq X, Tjian R. 2020. Overcoming the bottleneck to widespread testing: A rapid review of nucleic acid testing approaches for COVID-19 detection. RNA. http://dx.doi.org/10.1261/rna.076232.120.

Etievant S, Bal A, Escurret V, Brengel-Pesce K, Bouscambert M, Cheynet V, Generenaz L, Oriol G, Destras G, Billaud G, et al. 2020. Sensitivity assessment of SARS-CoV-2 PCR assays developed by WHO referral laboratories. medRxiv. http://dx.doi.org/10.1101/2020.05.03.20072207.

Kampf G, Todt D, Pfaender S, Steinmann E. 2020. Persistence of coronaviruses on inanimate surfaces and their inactivation with biocidal agents. Journal of Hospital Infection 104: 246–251.

Lemire KA, Rodriguez YY, McIntosh MT. 2016. Alkaline hydrolysis to remove potentially infectious viral RNA contaminants from DNA. Virol J 13: 88.

Leung RLC, Robinson MDM, Ajabali AAA, Karunanithy G, Lyons B, Raj R, Raoufmoghaddam S, Mohammed S, Claridge TDW, Baldwin AJ, et al. 2017. Monitoring the Disassembly of Virus-like Particles by 19F-NMR. Journal of the American Chemical Society 139: 5277–5280.

Li L, He J-A, Wang W, Xia Y, Song L, Chen Z-H, Zuo H-Z, Tan X-P, Ho AH-P, Kong S-K, et al. 2019. Development of a direct reverse-transcription quantitative PCR (dirRT-qPCR) assay for clinical Zika diagnosis. Int J Infect Dis 85: 167–174.

Liu Y, Yan L-M, Wan L, Xiang T-X, Le A, Liu J-M, Peiris M, Poon LLM, Zhang W. 2020. Viral dynamics in mild and severe cases of COVID-19. Lancet Infect Dis. http://dx.doi.org/10.1016/S1473-3099(20)30232-2.

McIntyre A, Patiar S, Wigfield S, Li J-L, Ledaki I, Turley H, Leek R, Snell C, Gatter K, Sly WS, et al. 2012. Carbonic anhydrase IX promotes tumor growth and necrosis in vivo and inhibition enhances anti-VEGF therapy. Clin Cancer Res 18: 3100–3111.

Nicola M, Alsafi Z, Sohrabi C, Kerwan A, Al-Jabir A, Iosifidis C, Agha M, Agha R. 2020. The Socio-Economic Implications of the Coronavirus and COVID-19 Pandemic: A Review. Int J Surg. http://dx.doi.org/10.1016/j.ijsu.2020.04.018.

Pandit P, Cooper-White J, Punyadeera C. 2013. High-yield RNA-extraction Method for Saliva. Clin Chem 59. http://dx.doi.org/10.1373/clinchem.2012.197863.

Pan Y, Long L, Zhang D, Yuan T, Cui S, Yang P, Wang Q, Ren S. 2020. Potential False-Negative Nucleic Acid Testing Results for Severe Acute Respiratory Syndrome Coronavirus 2 from Thermal Inactivation of Samples with Low Viral Loads. Clin Chem. http://dx.doi.org/10.1093/clinchem/hvaa091.

Pfaender S, Brinkmann J, Todt D, Riebesehl N, Steinmann J, Steinmann J, Pietschmann T, Steinmann E. 2015. Mechanisms of methods for hepatitis C virus inactivation. Appl Environ Microbiol 81: 1616–1621.

Prather KA, Wang CC, Schooley RT. 2020. Reducing transmission of SARS-CoV-2. Science eabc6197.

Rahman H, Carter I, Basile K, Donovan L, Kumar S, Tran T, Ko D, Alderson S, Sivaruban T, Eden J-S, et al. 2020. Interpret with caution: An evaluation of the commercial AusDiagnostics versus in-house developed assays for the detection of SARS-CoV-2 virus. J Clin Virol 127: 104374.

Reineke K, Mathys A, Knorr D. 2011. Shift of pH-Value During Thermal Treatments in Buffer Solutions and Selected Foods. International Journal of Food Properties 14: 870–881.

Roberts PL, Lloyd D, Marshall PJ. 2009. Virus inactivation in a factor VIII/VWF concentrate treated using a solvent/detergent procedure based on polysorbate 20. Biologicals 37: 26–31.

Rodino KG, Espy MJ, Buckwalter SP, Walchak RC, Germer JJ, Fernholz E, Boerger A, Schuetz AN, Yao JD, Binnicker MJ. 2020. Evaluation of saline, phosphate buffered saline and minimum essential medium as potential alternatives to viral transport media for SARS-CoV-2 testing. J Clin Microbiol. http://dx.doi.org/10.1128/JCM.00590-20.

Ruijter JM, Ramakers C, Hoogaars WMH, Karlen Y, Bakker O, van den Hoff MJB, Moorman AFM. 2009. Amplification efficiency: linking baseline and bias in the analysis of quantitative PCR data. Nucleic Acids Res 37: e45.

Satyanarayana M. 2020. Faulty probes are to blame for CDC coronavirus testing woes. Chemical and Engineering News https://cen.acs.org/analytical-chemistry/diagnostics/Faulty-probes-blame-CDC-coronavirus/98/i9.

Sentmanat M, Kouranova E, Cui X. 2020. One-step RNA extraction for RT-qPCR detection of 2019-nCoV. Molecular Biology 514.

Sullivan R, Heavey S, Graham DG, Wellman R, Khan S, Thrumurthy S, Simpson BS, Baker T, Jevons S, Ariza J, et al. 2020. An optimised saliva collection method to produce high-yield, high-quality RNA for translational research. PLoS One 15: e0229791.

To KK-W, Tsang OT-Y, Leung W-S, Tam AR, Wu T-C, Lung DC, Yip CC-Y, Cai J-P, Chan JM-C, Chik TS-H, et al. 2020a. Temporal profiles of viral load in posterior oropharyngeal saliva samples and serum antibody responses during infection by SARS-CoV-2: an observational cohort study. Lancet Infect Dis 20: 565–574.

To KK-W, Tsang OT-Y, Yip CC-Y, Chan K-H, Wu T-C, Chan JM-C, Leung W-S, Chik TS-H, Choi CY-C, Kandamby DH, et al. 2020b. Consistent Detection of 2019 Novel Coronavirus in Saliva. Clin Infect Dis. http://dx.doi.org/10.1093/cid/ciaa149.

UNSDG. 2020. SHARED RESPONSIBILITY, GLOBAL SOLIDARITY: Responding to the socio-economic impacts of COVID-19. United Nations https://unsdg.un.org/resources/shared-responsibility-global-solidarity-responding-socio-economic-impacts-covid-19.

Vogels CBF, Brito AF, Wyllie AL, Fauver JR, Ott IM, Kalinich CC, Petrone ME, Casanovas-Massana A, Catherine Muenker M, Moore AJ, et al. 2020. Analytical sensitivity and efficiency comparisons of SARS-COV-2 qRT-PCR primer-probe sets. medRxiv 2020.03.30.20048108.

WHO. 2020a. Coronavirus Disease 2019 (COVID-19) Situation Report – 125. World Health Organization https://www.who.int/docs/default-source/coronaviruse/situation-reports/20200524-covid-19-sitrep-125.pdf?sfvrsn=80e7d7f0_2.

WHO. 2020b. nCoV-2019 PCR protocol. World Health Organisation https://www.who.int/docs/default-source/coronaviruse/whoinhouseassays.pdf?sfvrsn=de3a76aa_2.

Xu R, Cui B, Duan X, Zhang P, Zhou X, Yuan Q. 2020. Saliva: potential diagnostic value and transmission of 2019-nCoV. Int J Oral Sci 12: 1–6.

Yasukawa K, Konishi A, Inouye K. 2010. Effects of organic solvents on the reverse transcription reaction catalyzed by reverse transcriptases from avian myeloblastosis virus and Moloney murine leukemia virus. Biosci Biotechnol Biochem 74: 1925–1930.

Ye J, Coulouris G, Zaretskaya I, Cutcutache I, Rozen S, Madden TL. 2012. Primer-BLAST: a tool to design target-specific primers for polymerase chain reaction. BMC Bioinformatics 13: 134.

